# Optogenetic inhibition of colon epithelium reduces hypersensitivity in a mouse model of inflammatory bowel disease

**DOI:** 10.1101/2020.05.21.107110

**Authors:** Sarah A. Najjar, Lindsay L. Ejoh, Emanuel Loeza-Alcocer, Brian S. Edwards, Kristen M. Smith-Edwards, Ariel Y. Epouhe, Michael S. Gold, Brian M. Davis, Kathryn M. Albers

**Affiliations:** Department of Neurobiology, University of Pittsburgh School of Medicine, Pittsburgh, Pennsylvania; Pittsburgh Center for Pain Research, University of Pittsburgh, Pennsylvania; Center for Neuroscience at the University of Pittsburgh, Pittsburgh, Pennsylvania

**Author notes:** For correspondence: Kathryn M. Albers, University of Pittsburgh, Department of Neurobiology, 200 Lothrop Street, Pittsburgh, PA 15216.

## Abstract

Visceral pain is a prevalent symptom of inflammatory bowel disease (IBD) that can be difficult to treat. Pain and hypersensitivity are mediated by extrinsic primary afferent neurons (ExPANs) that innervate the colon. Recent studies indicate that the colon epithelium contributes to initiating ExPAN firing and nociceptive responses. Based on these findings we hypothesized that the epithelium contributes to inflammation-induced hypersensitivity. A key prediction of this hypothesis is that inhibition of the epithelium would attenuate nociceptive signaling and inflammatory hypersensitivity. To test this hypothesis, the inhibitory yellow light activated protein archaerhodopsin was targeted to the intestinal epithelium (villin-Arch) or the ExPANs (TRPV1-Arch) that innervate the colon. Visceral sensitivity was assessed by measuring the visceromotor response (VMR) to colorectal distension (CRD), with and without yellow light illumination of the colon lumen. Inhibition of the colon epithelium in healthy villin-Arch mice significantly diminished the CRD-induced VMR. Direct inhibition of ExPANs during CRD using TRPV1-Arch mice showed that ExPAN and epithelial inhibition were similarly effective in reducing the VMR to CRD. We then investigated the effect of epithelial and ExPAN inhibition in the dextran sulfate sodium (DSS) model of inflammatory bowel disease (IBD). Inhibition of the colon epithelium significantly decreased DSS-induced hypersensitivity and was comparable to inhibition of ExPANS. Together these results reveal the potential of targeting the colon epithelium for treatment of pain.

## INTRODUCTION

In the gut, extrinsic primary afferent neurons (ExPANs) convey mechanical and chemical stimuli to neurons in the central nervous system [7]. In response to inflammatory and other pathophysiological changes in the gut, sensory afferents can become sensitized. Evidence suggests this sensitization significantly contributes to the chronic pain and decreased quality of life that accompany inflammatory bowel disease conditions [IBD; e.g., Crohn’s disease (CD) and ulcerative colitis (UC)] [6,12,23,24].

While there is evidence to suggest that intrinsic changes in ExPANs underlie the increase in excitability observed following inflammation [2,13-15], a growing body of evidence suggests changes in other cell types may also contribute to this process. Colon epithelial cells are one of these cell types. The epithelium not only maintains the mucosal barrier, but also synthesizes and releases neurotransmitters such as adenosine triphosphate (ATP) and 5-hydroxytrypamine (5-HT/serotonin) [26]. The spatially- and temporally-controlled release of these neurotransmitters is thought to facilitate communication between the epithelial lining and ExPAN terminals [5,20], supporting an important role for the epithelium in regulating primary afferent activity and nociceptive responses [3,4,22,25,27,31].

In the context of IBD, most studies of epithelial cells have focused on a failure of the epithelium to maintain a barrier, enabling bacterial infiltration and ExPAN sensitization secondary to the resulting immune response [28,29]. However, with evidence of a direct role in pain signaling, it is possible that the epithelium contributes to persistent ExPAN sensitization. Studies in animal models as well as in colon tissue from IBD patients suggest that inflammation affects enteroendocrine cell signaling [21], thus it is possible that inflammation changes normal communication between the epithelium and ExPANs. This may contribute to the sensitization of ExPANs and consequently the pain and hypersensitivity associated with IBD. To begin to address this possibility we tested whether optogenetic-mediated inhibition of the epithelium alters nociceptive signaling. We examined epithelial-sensory afferent communication under normal and inflamed conditions using a mouse model in which the inhibitory, yellow light activated archaerhodopsin (Arch) protein was targeted to epithelial cells. Colon sensitivity was assessed using the visceromotor response (VMR) to colorectal distension (CRD) measured with and without the light-induced inhibition of epithelial cells. We measured the efficacy of Arch-mediated inhibition of epithelial cells in reducing the VMR to CRD and then compared these results to those obtained in parallel using mice that express Arch in transient receptor potential vanilloid 1 (TRPV1)-lineage ExPANs. Results show that inhibition of the colon epithelium is effective in reducing inflammation-induced hypersensitivity.

## METHODS

### Animals

Male and female mice aged 6-8 months were analyzed. Two mouse lines were used: Vil-Arch, in which the inhibitory opsin ArchT was targeted to intestinal epithelial cells, and TRPV1-Arch, in which ArchT was targeted to primary afferent neurons innervating the colon. ArchT, an archaerhodopsin from *Halorubrum* strain TP009, is a yellow light driven outward proton pump that causes cellular hyperpolarization [17].

Mice with an ArchT-EGFP fusion protein in the *Rosa26* locus (Ai40D mice; RRID: IMSR_JAX:021188) were crossed with villin-Cre mice (RRID: IMSR_JAX:004586) or TRPV1-Cre mice (RRID:IMSR_JAX:017769). Littermates with Cre only or ArchT-EGFP only were used as controls. Animals were housed in an AAALAC-approved facility and handled in accordance with protocols approved by the University of Pittsburgh Institutional Animal Care and Use Committee.

### Laser-balloon device construction

To enable balloon distension simultaneous with optogenetic stimulation a 100-cm optical fiber (400 µm core with 1.25 mm ceramic ferrule and 440 µm FC/PC connector; Thor Labs) was threaded through a 15-cm piece of PE-200 polyethylene tubing connected to a 3-way stopcock [22]. To make the balloon, a 3 cm x 3 cm piece of Saran wrap was secured around the tubing using silk sutures (Ethicon). The balloon length was 1.5 cm and when inflated, its diameter was 0.9 cm. The fiber optic was attached to a 589 nm laser and power source (Laserglow Technologies). The laser power measured when illuminated through the plastic balloon was 12 mW/mm^2^.

### Electromyographic recording of visceromotor responses

Mice were fasted overnight and anesthetized using an intraperitoneal injection of urethane (Sigma; 1.2 g/kg). Mice were then placed on a heating pad and into a nosecone attached to an isoflurane vaporizer delivering 1% isoflurane (vaporized with 95% O_2_/5% CO_2_) (Parkland Scientific). The lower left quadrant of the abdomen was shaved, and a 1.5-cm incision was made in the skin, revealing abdominal musculature. Two clip electrodes (Pomona Electronics) were attached to the abdominal muscle, approximately 3 mm apart, to enable electromyographic (EMG) recording. The laser-balloon device was inserted into the colorectum and secured with tape to the tail. A ground electrode was secured with SignaGel (Parker Laboratories) to the mouse’s tail. Electrodes were attached to a differential amplifier (A-M Systems) connected to an A-D converter (Cambridge Electronic Design (CED) Micro 1401). EMG signals were amplified (10,000x), filtered (0.3-10 kHz band pass), and sampled at 20 kHz with the 1401 interface. Signals were recorded using Spike2 software (CED) and saved to a PC. Colorectal distension (CRD) was performed by balloon inflation using a compressed N_2_ tank equipped with a pressure regulator and connected to an air valve box enabling computer control. Prior to testing isoflurane level was slowly lowered to 0.25%. Mice were held at this level of combined anesthesia (isoflurane with previous urethane injection) until they showed no signs of ambulation but were responsive to toe pinch. After the mice displayed consistent responses to 60 mmHg CRD, the stimulus protocol was administered. This consisted of 3-5 trials of 10 s 60 mmHg CRD, followed by 3-5 trials of 10 s 60 mmHg CRD plus yellow laser; laser was turned on 1 s before the distension stimulus and remained on during the 10 s stimulus. All stimuli were delivered at 4-min intervals. EMG waveforms were rectified in Spike2 and the integral of the waveform was used to quantify VMRs. The resting activity 10 s before distension was subtracted from the distension-evoked activity. EMG waveform extraction and data analysis were performed in a blinded manner.

### Dextran sulfate sodium (DSS) inflammation protocol

DSS was administered in drinking water. A 3% DSS solution was prepared by dissolving DSS (36,000-50,000 MW; MP Biomedicals) into autoclaved water that was provided *ad libitum* via water bottle for 5 days. Vehicle control mice were provided water without DSS. VMR analysis was performed immediately after 5 days of DSS treatment.

### Fluorescent microscopy and histopathological scoring

Distal colon segments and L6 dorsal root ganglia (DRG) were isolated and fixed in 4% paraformaldehyde for at least 30 min. Tissues were cryopreserved in 25% sucrose overnight, embedded in optimal cutting temperature (OCT) compound, and sectioned at 14 µm thickness. Sections were coverslipped and EGFP expression imaged using a Leica DM4000B microscope equipped with a Leica DFC7000 T digital camera.

DSS- and vehicle-treated mice were euthanized with isoflurane after VMR experiments. A 1-cm piece of colon was removed and fixed for at least 30 min in 4% paraformaldehyde in 0.01 M phosphate-buffered saline (PBS) and cryoprotected in 25% sucrose in PBS at 4°C for 24 hours. The tissue was embedded in OCT, sectioned on a cryostat at 14 μm, and mounted on Superfrost Plus microscope slides (Fisherbrand). Sections were stained with hematoxylin and eosin (H&E) and brightfield images captured using a Leica DM4000B microscope with a digital camera attachment.

For disease scoring, a modified method of histopathological analysis was used [19]. The extent of damage to the tissue was determined by goblet cell depletion (0 = absent, 2 = severe), inflammatory cell infiltration (0 = normal, 2 = dense) and thickening of the submucosal layer (0 = base of crypt sits on muscularis mucosa, 2 = marked muscle thickening present). The disease score is the sum of the scores from each of these three categories, with a max score of 6 signifying greater pathological damage to the colon tissue. The experimenter determining disease scores was blinded to the treatment mice received (DSS vs. vehicle).

### Data analysis

Statistical analyses were performed using Prism (GraphPad Software, San Diego, CA). For VMR experiments, mice received 3-5 trials of 60 mmHg CRD to establish baseline responses and then 3-5 trials of 60 mmHg CRD + Laser. Trials with laser were considered inhibited when there was a significant (>2 standard deviations) decrease in VMR from mean baseline responses. Comparisons between Vil-Arch and TRPV1-Arch mice that displayed yellow-light induced inhibition in VMR (i.e., “responders”) were performed using 2-way ANOVA and Mann-Whitney tests. Histopathological scores and spleen weights of the DSS-treated colons were compared to vehicle-treated colons using unpaired t-tests. VMR to CRD at various pressures was tested in DSS-treated and untreated mice using a two-way ANOVA. Statistical tests are specified in the results section and significance was defined as p < 0.05. Data are plotted as mean ± standard error of the mean.

## RESULTS

### Optogenetic inhibition of colon epithelial cells reduces visceromotor responses to colorectal distension

To enable specific inhibition of the colon epithelium, the yellow light-activated proton pump Arch was expressed under control of the villin-Cre driver. In Vil-Arch animals, Arch is fused to enhanced green fluorescent protein (EGFP), enabling visualization of the fusion protein in the villin-expressing lining epithelial cells of the colon (**Fig. 1A**). Arch-EGFP expression was restricted to the colon epithelium and not present in the L6 DRG (**Fig. 1B**). A laser-balloon device was constructed to enable colorectal distension (CRD) simultaneous with light illumination of the colon lumen (**Fig. 1C**). Visceromotor responses (VMR) to CRD were recorded, as they provide a reliable indication of visceral nociception [11]. VMRs to noxious distension (3-5 trials of 60 mmHg CRD) were recorded in Vil-Arch and control littermate mice before (baseline) and during yellow light application (**Fig. 1D**). Of 7 Vil-Arch mice tested, 4 displayed inhibition with yellow light. Inhibition was defined as VMR <2 standard deviations from average baseline responses. In the CRD trials that were inhibited, yellow light reduced the VMR by 67 ± 19% (VMR to CRD baseline: 2.67 ± 0.72 V*s vs. VMR to CRD + laser: 1.14 ± 0.88 V*s; **Fig. 1E**). This inhibition occurred on average in 66% of CRD + laser trials; **Fig. 1E inset**). The remaining 3 Vil-Arch mice that were tested did not exhibit a decrease in VMR in response to light stimulation that met our threshold of <2 standard deviations from average baseline responses (VMR to CRD baseline: 1.4 ± 0.52 V*s vs. average VMR to CRD + laser: 1.39 ± 0.60 V*s; **Fig. 1F**). The littermate control mice (n = 5) did not display inhibition with yellow light in any trial (VMR to CRD baseline: 1.65 ± 0.27 V*s vs. average VMR to CRD + laser: 1.71 ± 0.32 V*s; **Fig. 1G**).

**Figure 1.**
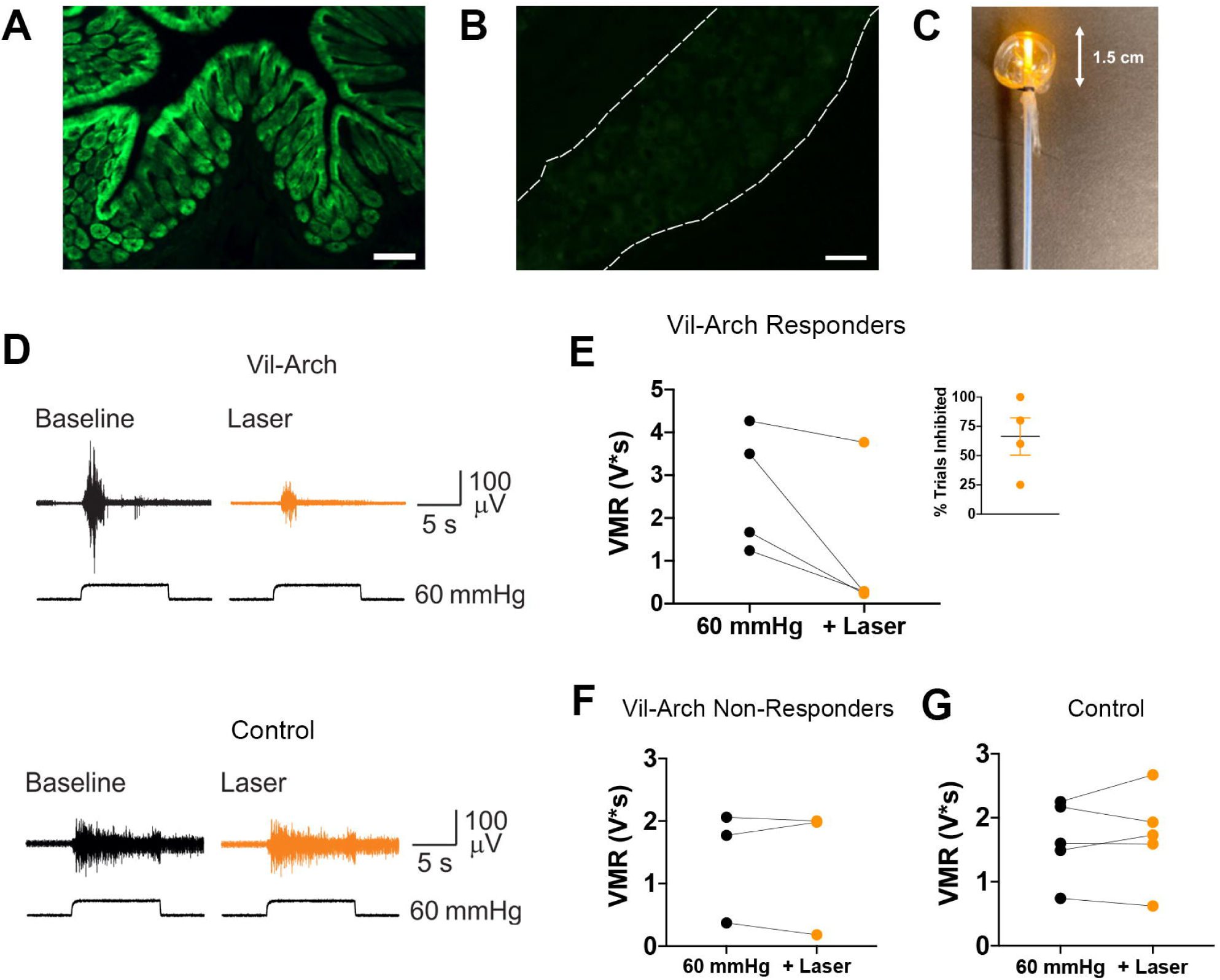
Inhibition of the colon epithelium reduces visceromotor response (VMR) to colorectal distension (CRD). Inhibitory opsin archaerhodopsin (Arch) is conjugated to enhanced green fluorescent protein (EGFP) to enable visualization. Arch-EGFP (green) is specifically expressed in colon epithelial cells under the villin-cre driver **(A)** and not in the L6 DRG **(B). C)** A laser-balloon device was constructed to enable CRD simultaneous with light illumination of the colon lumen. **D)** VMRs to 60 mmHg CRD were recorded in villin-Archaerhodopsin (Vil-Arch) and control littermate mice before (Baseline) and during yellow light application (+ Laser). **E)** Out of 7 Vil-Arch mice, 4 displayed a significant decrease in VMR to CRD with the addition of yellow light, which occurred in an average of 51% of trials (**inset**). **F)** In the remaining 3 Vil-Arch mice that were tested, none of the CRD + Laser trials met the threshold for a decrease. **G)** Littermate control mice (n = 5) did not display inhibition with yellow light in any trial. Scale bars = 100 *µ*M (A), 50 *µ*M (B).

### Optogenetic inhibition of colon ExPANs reduces VMR to CRD

For comparison, we next measured changes in distension-induced VMRs due to yellow light stimulation in TRPV1-Arch mice. In these experiments, Arch was targeted to colon extrinsic primary afferent neurons (ExPANs) using the TRPV1-Cre driver. As TRPV1 is expressed in nearly 90% of colon afferents [23], Arch-EGFP expression was observed in nerve terminals in the colon (**Fig. 2A**) and in cell bodies in the DRG (**Fig. 2B**). VMRs to 60 mmHg CRD were recorded in TRPV1-Arch and control littermate mice before and during yellow light stimulation (**Fig. 2C**). The same criteria were employed to identify mice responsive to light stimulation. Of 6 TRPV1-Arch mice tested, 4 responded to yellow light stimulation. In the trials in which CRD was inhibited, yellow light reduced the VMR by 84 ± 8% (VMR to CRD baseline: 1.22 ± 0.12 V*s vs. VMR to CRD + laser: 0.18 ± 0.08 V*s; **Fig. 2D**). This inhibition occurred on average in 82% of trials (**Fig. 2D inset**). The remaining 2 TRPV1-Arch mice did not respond to yellow light (**Fig. 2E**). In control littermate mice (n = 4), no reduction in VMR in response to light stimulation was observed (baseline VMR to CRD: 1.84 ± 0.23 V*s vs. average VMR to CRD + laser: 1.82 ± 0.33 V*s; **Fig. 2F**). When comparing the Vil-Arch and TRPV1-Arch mice that responded to yellow light, there was no statistical difference between groups in either the extent of inhibition in which a response was detected [main effect of yellow light (p = 0.01), no effect of genotype (p = 0.16), 2-way ANOVA; **Fig. 2G**] or in the percentage of trials in which inhibition was detected (p = 0.57, Mann-Whitney test; **Fig. 2H**).

**Figure 2.**
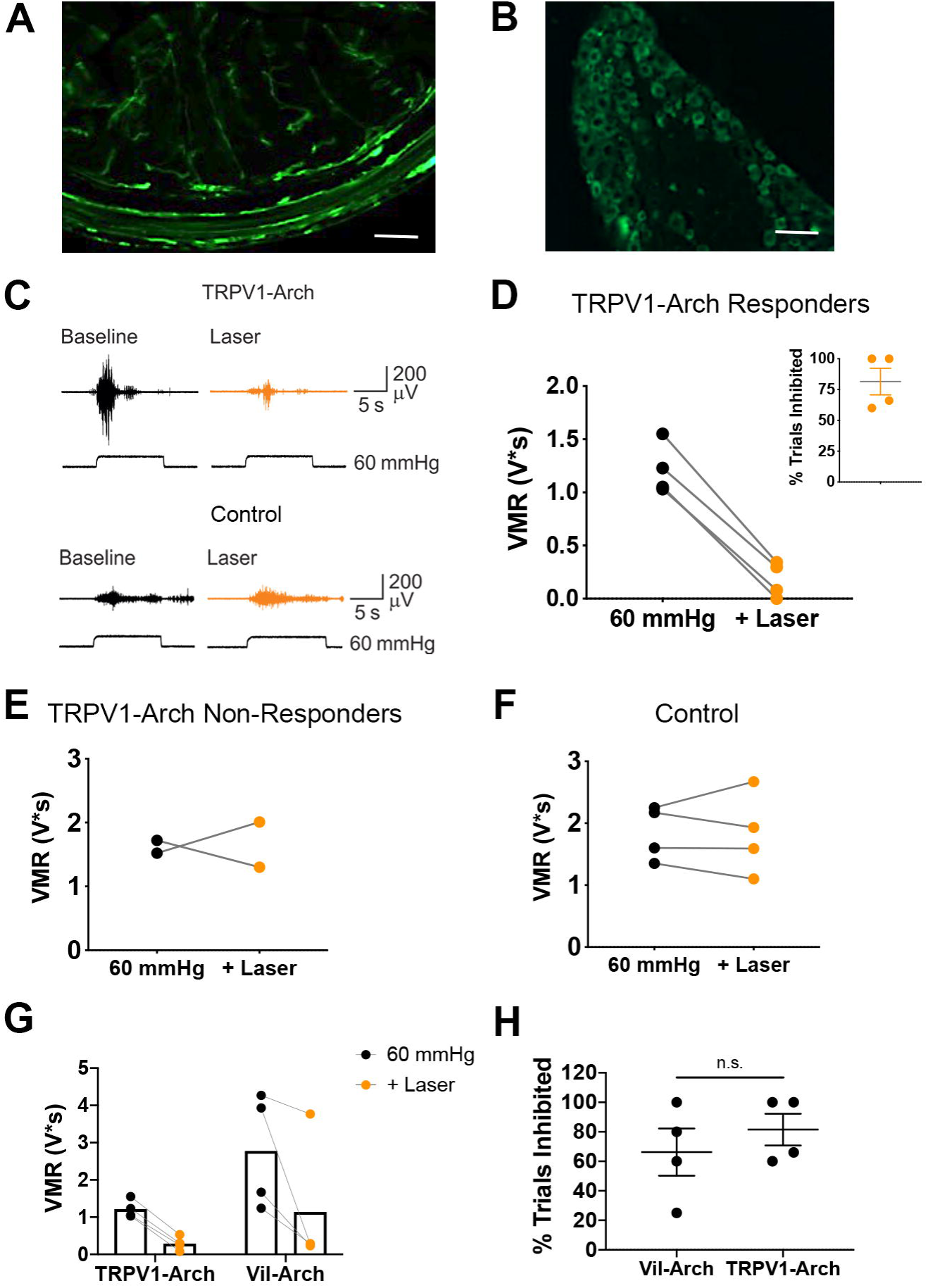
Inhibition of colon extrinsic primary afferent neurons reduces VMRs to colorectal distention. **A)** Arch-EGFP was targeted to colon afferents using the TRPV1-cre driver. Arch-EGFP-positive nerve terminals innervate the distal colon. **B)** Arch-EGFP-positive (green) nerve cell bodies are present in the L6 DRG. **C)** VMRs to 60 mmHg CRD were recorded in TRPV1-Arch and control littermate mice before (Baseline) and during yellow light illumination (+ Laser). **D)** Out of 6 TRPV1-Arch mice, 4 displayed a significant decrease in VMR to CRD with the addition of yellow light, which occurred in an average of 82% of trials (**inset**). **E)** The remaining 2 TRPV1-Arch mice did not display yellow light-induced reduction in VMR. **F)** In the control littermate mice (n = 4), no reduction in VMR in response to yellow light was observed. **G)** In comparing the Vil-Arch and TRPV1-Arch mice that responded to yellow light (n = 4 per group), no statistical differences were detected in the extent of inhibition in responding trials [main effect of yellow light (p = 0.01), no effect of genotype (p = 0.16), 2-way ANOVA]. **H)** There was also no significant difference in the percentage of inhibited trials in Vil-Arch vs. TRPV1-Arch responders (p = 0.57, Mann-Whitney test).

### DSS-mediated inflammation causes visceral hypersensitivity

To examine the role of epithelial cells in inflammation-induced hypersensitivity, we used the dextran sulfate sodium (DSS) model of inflammatory bowel disease. DSS damages the mucosal barrier, causing inflammation and other symptoms that mimic human ulcerative colitis [9]. After testing dose and time variables, we used a protocol in which 3% DSS is administered in the drinking water for 5 days prior to VMR analysis, as it consistently produced hypersensitivity. Histopathological comparison of vehicle-treated (**Fig. 3A**) versus DSS-treated (**Fig. 3B**) colons showed that this DSS protocol did not cause notable loss of lining epithelial cells, which allowed further examination of epithelial-neuronal communication. In addition, fluorescent images of vehicle-treated (**Fig. 3C**) versus DSS-treated (**Fig. 3D**) Vil-Arch colons showed that Arch-EGFP expression was not reduced by DSS treatment.

**Figure 3.**
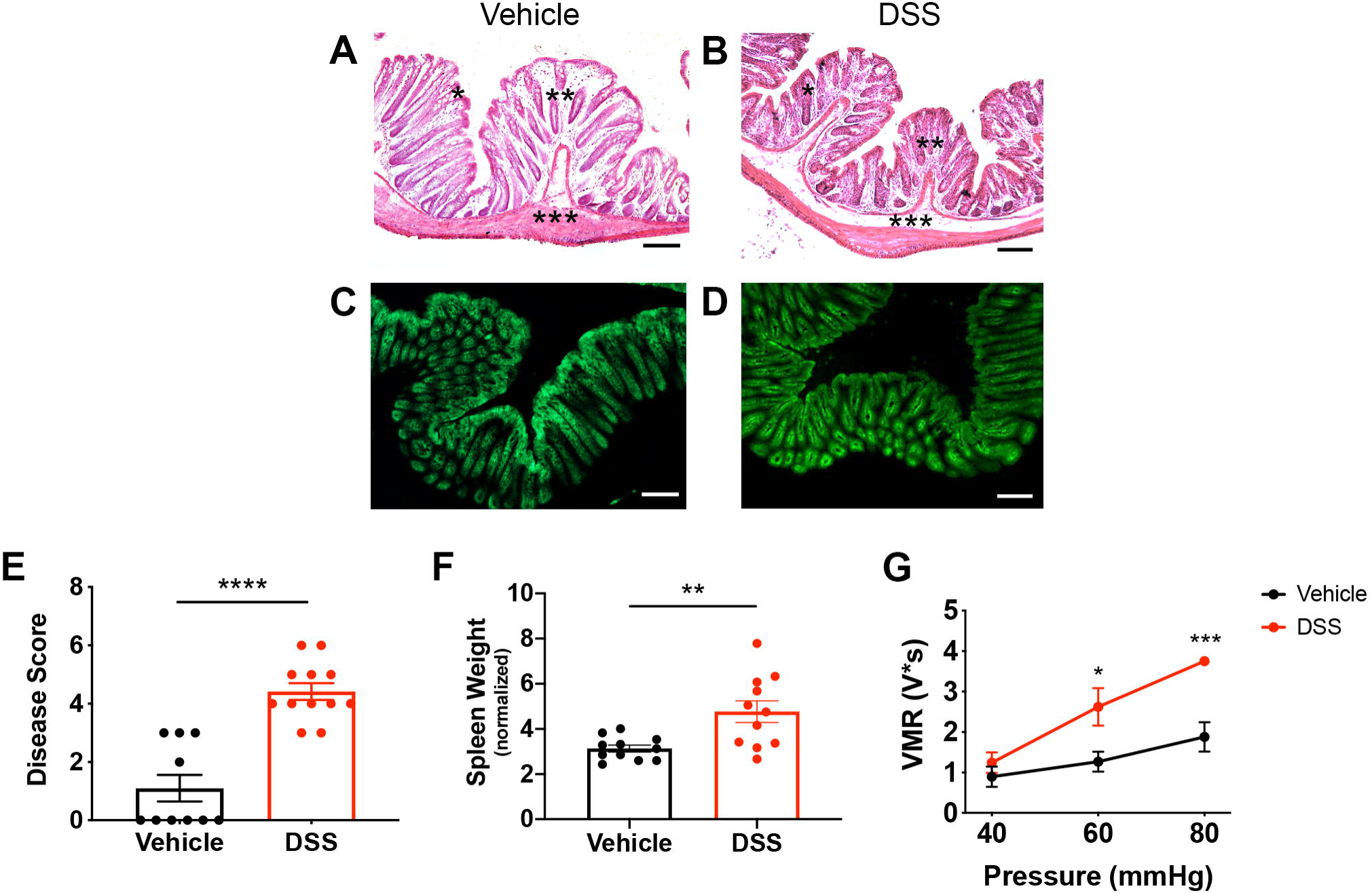
DSS treatment induces colon inflammation and visceral hypersensitivity. Mice were administered 3% DSS for 5 days or vehicle treatment. Histopathological analysis of vehicle-treated **(A)** vs. DSS-treated **(B)** colons showed that the DSS protocol did not cause a notable loss in lining epithelial cells. There was a depletion of goblet cells (indicated by *), increase in infiltrating cells (indicated by **), and thickening of the submucosal layer (indicated by ***). Fluorescent images of vehicle-treated **(C)** and DSS-treated **(D)** Vil-Arch colons showed comparable EGFP expression, indicating that Arch-EGFP expression is not ablated after DSS treatment. **E)** DSS-treated mice had a higher inflammation disease score, indicating fewer mucous-secreting goblet cells, more infiltrating cells, and expansion of the submucosa (n = 10 in vehicle-treated group and n = 12 in DSS-treated group; p < 0.0001, unpaired t-test). **F)** DSS-treated mice also had significantly higher spleen weights indicating inflammation n = 11 in vehicle-treated group and n = 11 in DSS-treated group; p = 0.007, unpaired t-test with Welch’s correction. **G)** Visceromotor responses (VMRs) to colorectal distension (CRD) were measured in vehicle- and DSS-treated mice. Mice received multiple CRD pressures (40, 60, and 80 mmHg) and showed significantly greater VMRs at 60 (p = 0.01) and 80 mmHg (p < 0.001, two-way ANOVA with Sidak’s test; n = 5 mice per group). Scale bars = 100 *µ*M.

Though the epithelial lining is intact, histopathological analysis [19] showed that DSS-treated mice had a higher inflammation score, as indicated by fewer mucous-secreting goblet cells, more infiltrating immune cells, and thickening of the submucosal layer ((n = 10 in vehicle-treated group and n = 12 in DSS-treated group; p < 0001, unpaired t-test; **Fig. 3E**). Increased spleen weights in DSS-treated mice, an indicator of inflammation, were also present (n = 11 in vehicle-treated group and n = 11 in DSS-treated group; p = 0.007, unpaired t-test with Welch’s correction; **Fig. 3F**). These data were pooled from vehicle- and DSS-treated Vil-Arch, TRPV1-Arch and control mice, as previous analysis showed that genotype did not have an effect on inflammation. When comparing disease scores in DSS-treated Vil-Arch, TRPV1-Arch and control mice, there was a main effect of DSS (p < 0.0001) but not genotype (p = 0.09), and there was no interaction between DSS and genotype (p = 0.16; two-way ANOVA). When comparing spleen weights, there was a main effect of treatment (p = 0.01) but not genotype (p = 0.91), and there was no interaction (p = 0.99; two-way ANOVA).

Previous studies have shown that DSS treatment using various dose protocols can result in visceral hypersensitivity [18,30,32]. To confirm that our DSS protocol produced hypersensitivity, VMR analysis was conducted in Vil-Arch mice. Mice were stimulated with CRD pressures of 40, 60, and 80 mmHg. There were main effects of DSS treatment (p = 0.02) and distension pressure (p < 0.0001) with a significant interaction (p = 0.002, two-way ANOVA). In addition, the VMR in response to 60 and 80 mmHg was significantly greater in DSS treated than vehicle-treated mice (p = 0.01 and p < 0.001, respectively, Sidak’s test; n = 5 mice per group) (**Fig. 3G**).

### Inhibition of the colon epithelium via Arch-activation reduces DSS-induced hypersensitivity

As our DSS protocol produced hypersensitivity to CRD in response to 60 mmHg pressure, we used this stimulus intensity to assess the impact of yellow light stimulation on the VMR in DSS-treated control and Vil-Arch mice. VMRs to CRD were recorded in Vil-Arch and control littermate mice, both treated with DSS. Measures were made before (baseline) and during application of light stimulation (**Fig. 4A**). Yellow light produced effective inhibition in 4 out of 5 Vil-Arch mice, reducing the VMR by 51 ± 9% (baseline VMR to CRD: 2.66 ± 0.57 V*s vs. VMR to CRD + laser: 1.22 ± 0.41 V*s; **Fig. 4B**). Yellow light was effective in inhibiting the VMR in 60 ± 9% of trials (**Fig. 4B inset**). In the 1 Vil-Arch mouse that did not respond to yellow light, the baseline VMR to CRD was 2.58 V*s and the average VMR to CRD + laser was 2.49 V*s. In the control littermate mice (n = 4), no reduction in VMR in response to light stimulation was observed (baseline VMR to CRD: 2.1 ± 0.2 V*s vs. average VMR to CRD + laser: 2.68 ± 0.42 V*s; **Fig. 4C**). Yellow light induced inhibition in the DSS-treated Vil-Arch mice to an extent comparable to untreated Vil-Arch mice (vehicle: 67 ± 19% inhibition vs. DSS: 51 ± 9%; p = 0.57, unpaired t-test; **Fig 4D**).

**Figure 4.**
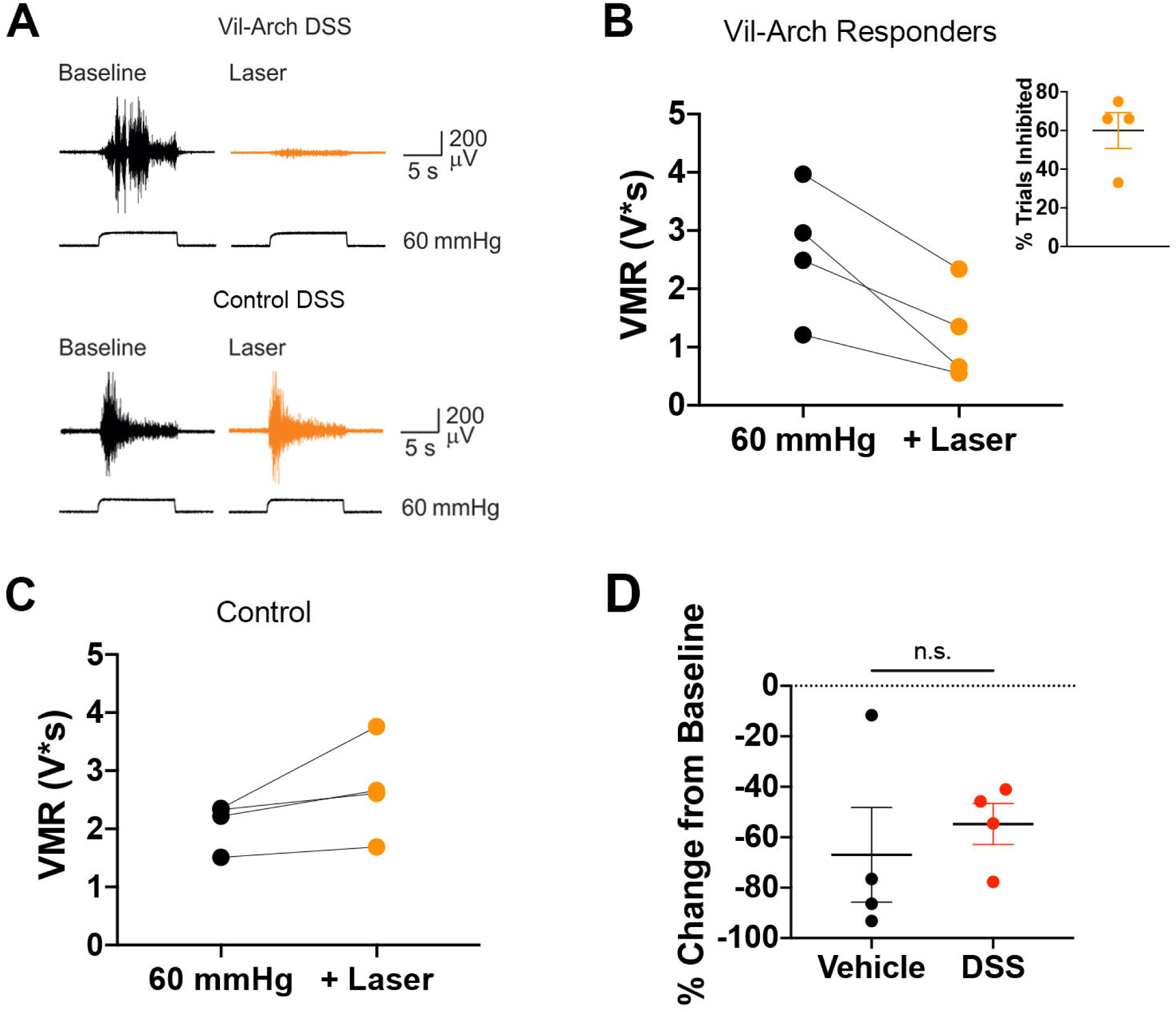
Inhibition of colon epithelium reduces DSS-induced hypersensitivity. **A)** VMRs to 60 mmHg colorectal distension were recorded in DSS-treated Vil-Arch and control littermate mice before (Baseline) and during yellow light stimulation (+ Laser). **B)** Yellow light produced effective inhibition in 4 out of 5 Vil-Arch mice, in an average of 60% of CRD + laser trials (**inset**). **C)** In control littermate mice (n = 4), no reduction in VMR in response to light stimulation was observed. **D)** Yellow light-induced inhibition in DSS-treated Vil-Arch mice was similar to inhibition in untreated Vil-Arch mice (n = 4 per group; p = 0.57, unpaired t-test).

### Inhibition of colon extrinsic primary afferent neurons reduces DSS-induced hypersensitivity

We next examined the effects of inhibiting colon afferents on DSS-induced hypersensitivity. VMRs to CRD were recorded in TRPV1-Arch and control littermate mice before (baseline) and during yellow light stimulation (**Fig. 5A**). Yellow light was effective in inhibiting the VMR to CRD in 5 out of 6 TRPV1-Arch mice. In the CRD trials that were inhibited, yellow light reduced the VMR by 62 ± 8% (baseline VMR to CRD: 2.17 ± 0.39 V*s vs. VMR to CRD + laser: 1.82 ± 0.31 V*s; **Fig. 5B**). Inhibition occurred in an average of 54% of the CRD + laser trials in these mice (**Fig. 5B inset**). In the 1 TRPV1-Arch mouse that did not respond to yellow light, the baseline VMR to CRD was 2.07 V*s and the average VMR to CRD + laser was 2.09 V*s. In control littermate mice (n = 4), yellow light-induced reduction in VMR only occurred in 1 CRD + laser trial in 1 mouse. In all other trials, this reduction was not observed (baseline VMR to CRD: 1.93 ± 0.45 V*s vs. average VMR to CRD + laser: 2.07 ± 0.39 V*s; **Fig. 5C**). Yellow light induced inhibition in the DSS-treated TRPV1-Arch was compared to inhibition in untreated TRPV1-Arch mice (vehicle: 83 ± 8% inhibition vs. DSS: 53 ± 11%; p = 0.11, Mann-Whitney test; **Fig 5D**). There was a trend towards a decrease in the extent of inhibition in DSS-treated mice, but this was not significant. Finally, comparison between inhibition via Arch-expression in the epithelium versus inhibition in colon afferents showed that both were similarly effective at reducing the VMR response in inflamed mice. No statistical differences were detected in the extent of inhibition in responding trials [main effect of yellow light (p = 0.001), no effect of genotype (p = 0.57), 2-way ANOVA; **Fig. 5E**] or in the rate of inhibited trials (p = 0.62, Mann-Whitney; **Fig. 5F**).

**Figure 5.**
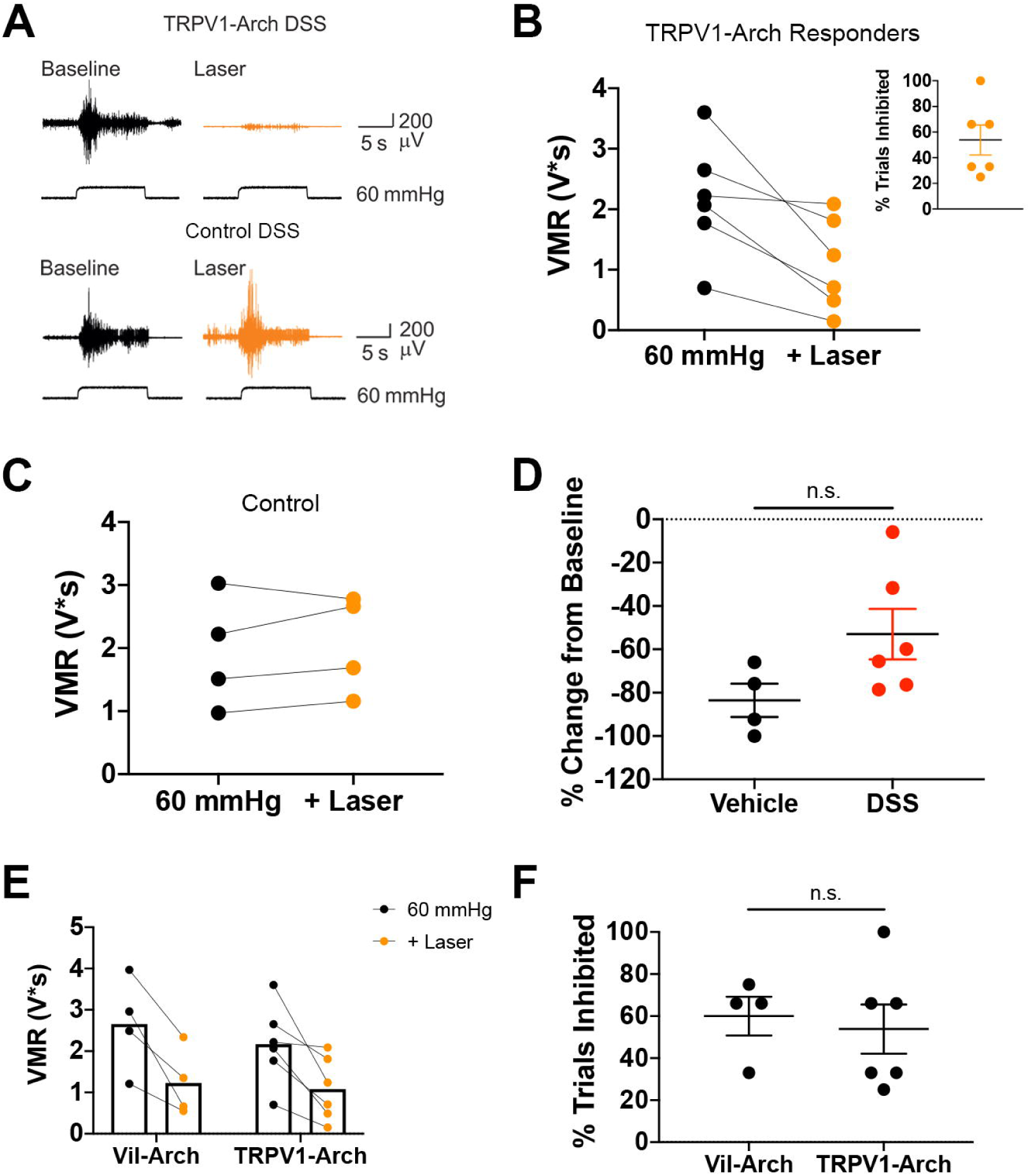
Inhibition of colon afferents reduces DSS-induced hypersensitivity. **A)** VMRs to colorectal distension were recorded in DSS-treated TRPV1-Arch and control littermate mice before (Baseline) and during yellow light stimulation (+ Laser). **B)** Yellow light produced effective inhibition in 5 out of 6 TRPV1-Arch mice, in an average of 45% of CRD + laser trials (**inset**). **C)** In control littermate mice (n = 4), no reduction in VMR in response to light stimulation was observed. **D)** Yellow light-induced inhibition in the DSS-treated TRPV1-Arch mice (n = 5) was compared to inhibition in untreated TRPV1-Arch mice (n = 4). There was a trend for a decrease in inhibition in DSS-treated mice, but this was not significant (p = 0.11; Mann-Whitney test). **E)** Comparing DSS-treated Vil-Arch (n = 4) and TRPV1-Arch (n = 5) mice that responded to yellow light shows no statistical differences in the extent of inhibition in responding trials [main effect of yellow light (p = 0.001), no effect of genotype (p = 0.57), 2-way ANOVA]. **F)** There was also no significant difference in the percentage of inhibited trials in Vil-Arch vs. TRPV1-Arch responders (p = 0.62, Mann-Whitney test).

## DISCUSSION

Arch-mediated inhibition of colon epithelial cells was found to diminish visceral nociceptive responses in both healthy mice and mice with DSS-induced inflammation. In Vil-Arch animals that displayed yellow light-induced decreases in VMR, inhibiting the epithelium reduced VMRs by more than 65%. Notably, epithelial inhibition was similarly effective as colon afferent inhibition in reducing the VMR. When combined with our previous findings that ChR2-mediated excitation of the colon epithelium evokes VMRs [22], these results support the role of the epithelium as a critical component of nociceptive signaling.

How changes in the epithelium specifically impact the response to noxious stimuli has proven difficult to study because of the complexity of neural and epithelial cell types and the intimate association of nerve terminals and the epithelium, e.g., chemical or mechanical stimulation of the colon simultaneously activates both nerves and epithelial cells, making the relative contribution of each cell type undecipherable. To overcome this limitation, we used villin-Cre driver mice to genetically target the inhibitory Arch opsin to all epithelial cells lining the colon. This allowed a specific assessment of the contribution of the epithelium to distension evoked VMRs and revealed that inhibition of the epithelium reduces visceral hypersensitivity. However, because the villin-Cre driver targets enterocyte, goblet, enterochromaffin and enteroendocrine cell subpopulations it is not possible to determine which of these subpopulations of epithelial cell types mediates the effects of light stimulation. Thus, future studies should employ additional Cre-lines to reveal the contribution of different epithelial cell types to visceral nociception.

Although VMR is a reliable way to assess visceral sensitivity [11], we observed variability in responses to CRD within the Vil-Arch and TRPV1-Arch groups. Some Arch-expressing mice did not display yellow light mediated inhibition of the VMR in any CRD + laser trials. This variability in VMR may be due to inappropriate positioning of the laser or excessive luminal contents that occluded the laser. The lack of a reliable response could also be due to the interstimulus interval used in these experiments. We consistently waited 4 min in between distension and distention + laser trials, and this may not have been enough time for the Arch proton pump to reset properly. Variability could also be attributed to the possibilities that different epithelial cell types play different roles in the regulation of ExPAN activity, or Arch activation has different effects in subpopulations of epithelial cells. That is, while previous studies showed light activation of Arch leads to proton release and membrane hyperpolarization, preventing action potential firing in neurons [10], it remains to be determined how Arch-mediated hyperpolarization of colon epithelial cell membranes results in the ‘silencing’ of epithelial cells and inhibition of distension-induced VMRs. One possibility is that hyperpolarization blocks the distension (stretch)-mediated release of ATP from the epithelium, which is known to activate purinergic receptors expressed on extrinsic afferent terminals [8,22]. Release of 5-HT from the subset of electrically excitable enterochromaffin cells, as shown in cultured organoids [1,4] may also be changed by Arch-mediated hyperpolarization.

Our studies indicate limitations in the extent to which inhibition of colon epithelial cells or ExPANs can reduce VMRs to CRD. Post DSS treatment, both epithelial and ExPAN inhibition were effective in reducing hypersensitivity. However, averaging of VMR values showed that neither could by itself completely *reverse* the hypersensitivity, i.e., blocking peripheral components of the VMR could not normalize sensitivity to CRD after inflammation. The inability to completely reverse hypersensitivity suggests that Arch-mediated inhibition did not sufficiently inhibit colon afferents. There could also be a role for central sensitization in this inflammatory state. Future studies will investigate how other components involved in the VMR circuit contribute to visceral hypersensitivity.

Visceral pain is notoriously difficult to treat, often persisting long after the precipitating injury or disease is no longer evident. Chronic disruptive changes in the epithelial lining may affect normal epithelial-neural signaling leading to persistence in hypersensitivity. Indeed, changes were identified in muscarinic signaling in urothelial cells isolated from the bladder of patients with chronic interstitial cystitis [16], suggesting injury can evoke long-term changes in epithelial signaling properties. It remains to be determined whether there are similar long-term changes in colon epithelial cells, although there is evidence that inflammation affects the abundance and neurotransmitter content of enteroendocrine cells [21]. The work here indicates the epithelium is an important component of nociceptive signaling and a potential target for treatment of visceral pain.

## ACKNOWLEDGEMENTS

We thank Mr. Christopher Sullivan for his excellent technical support in mouse colony maintenance and genetic screening. This work was supported by funding from the National Institutes of Health (T32 NS073548, T32 DK063922-17, OT2 OD023859, R01AR069951, R01 DK107966, and UG3 TR003090-01). All authors have no conflict of interest.

